# Gut microbiota and type 2 diabetes associaction: A meta-analysis of 16S studies and their methodological challenges

**DOI:** 10.1101/2024.10.01.615711

**Authors:** Jéssica Lígia Picanço Machado, Ana Paula Schaan, Izabela Mamede, Gabriel da Rocha Fernandes

## Abstract

Diabetes mellitus is a prevalent chronic non-communicable disease, and recent studies have explored the link between gut microbiota and its development. Despite some evidence suggesting an association, the influence of gut microbiota on type 2 diabetes DM2 remains unclear. A systematic search of PubMed Janeiro 2016– 10 December 2023 using the keywords “16S” and “diabetes” or “DM2” and “gut microbiota” and “diabetes” or “DM2”. The studies included compared gut microbiome diversity between diabetic and non-diabetic adults using 16S rRNA sequencing, excluding children, interventions, and type 1 diabetes. Alpha diversity indices and bacterial mean abundance were analyzed, with statistical assessments using a random-effects model and I^2^ for heterogeneity. Thirteen studies met the criteria, with the Shannon index being the most commonly used measure. Results showed significant heterogeneity I^2^ *>* 75% and no notable differences between diabetic and non-diabetic groups. Other indices, such as Chao1 and phylogenetic whole tree, similarly showed no consistent differences. Taxonomic analysis also failed to find phyla consistently correlated with DM2, with variability across studies. The relationship between gut microbiota and diabetes remains uncertain due to technical and biological factors that are often overlooked. The inconsistencies across studies highlight the low reproducibility common in microbiota research.

## 1 Introduction

As global populations increasingly adopt urbanized lifestyles, the prevalence of chronic noncommunicable diseases, such as diabetes mellitus DM, has become a significant public health concern, particularly in low- and middle-income countries (WHO 2023). It is estimated that over 500 million adults worldwide are living with DM, contributing to an enormous and growing economic burden (International Diabetes Federation 2021). Beyond its impact on glucose regulation, DM is a major risk factor for cardiovascular diseases, which remain the leading cause of death globally (WHO 2024).

DM, being a multifactorial group of diseases, is influenced by genetic predisposition, environmental factors, and has recently been linked to alterations in the gut microbiome (Gilbert *et al*., 2018; Qin *et al*., 2012). Numerous studies have proposed a role for the gut microbiome in the pathophysiology of DM, attributed to its influence on host metabolic homeostasis. This includes maintaining the gut epithelial barrier, maturing the immune system, and producing a variety of metabolites that exert systemic effects on the host (Vrieze *et al*., 2012; Rogers and Wesselingh 2016).

The relationship between the gut microbiota and DM, however, remains contentious, with inconsistent findings across different populations (He *et al*., 2018; Zhou *et al*., 2019). For instance, the genus Bacteroides has been reported to have both higher and lower relative abundance in diabetic patients across various studies (He *et al*., 2018; Yamaguchi *et al*., 2016). Some metaanalyses have highlighted this inconsistency, suggesting that the gut microbiome may not play a significant role in DM development (Gurung *et al*., 2020; MetaHIT consortium *et al*., 2015). This has led to the hypothesis that it is the overall functional repertoire and metabolic output of the microbial community, rather than specific taxa, that are critical in the interaction between the microbiome and DM (Vatanen *et al*., 2018).

Concerns about the reproducibility of metagenomic studies, particularly in methodology, have also emerged. Notably, a highly cited article foundational to many studies was found to have methodological flaws (Gihawi *et al*., 2023). In response to these issues, we conducted a meta-analysis of datasets where gut microbiota, assessed through 16S rRNA gene sequencing, was studied in relation to Diabetes Mellitus.

## 2 Method

This systematic review and meta-analysis aimed to evaluate the relationship between the gut microbiome and type 2 diabetes mellitus DM2 by analyzing 16S rRNA sequencing data. The study was designed to synthesize available evidence, identify patterns or discrepancies in the findings, and assess the reproducibility of results across different studies. Our approach followed the PRISMA Preferred Reporting Items for Systematic Reviews and Meta-Analyses guidelines to ensure a rigorous and transparent methodology.

### 2.1 Data Search

A comprehensive and systematic search was conducted in PubMed to identify relevant studies published between January 2016 to 14 December 2023. The search strategy combined Medical Subject Headings MeSH terms and keywords to capture all pertinent literature. The search string included the following terms: “16S” AND “diabetes” OR “DM2” AND “gut microbiota” AND “diabetes” OR “DM2”. To ensure the quality and relevance of the data, only peer-reviewed articles published in English were considered. The search was complemented by manual screening of reference lists from selected studies to identify any additional relevant publications.

### 2.2 Study Selection

The selection process involved a multi-step approach. Initially, titles and abstracts were screened to eliminate studies that clearly did not meet the inclusion criteria. Full-text reviews were then conducted for studies that appeared potentially eligible. Studies were included if they met the following criteria: 1 compared gut microbiome diversity between adult diabetic and non-diabetic populations; 2 employed 16S rRNA sequencing as the primary method for microbiome analysis; and 3 were published in English. Studies were excluded based on the following criteria: 1 studies involving pediatric populations, due to differences in microbiome composition; 2 studies relying solely on quantitative PCR qPCR for bacterial abundance, as this method lacks the depth of 16S rRNA sequencing; 3 studies employing shotgun sequencing, which differ significantly in methodology and scope from 16S studies; and 4 studies focusing primarily on inflammatory markers or other non-microbiome-related associations with diabetes. This rigorous selection process ensured that the included studies were comparable and relevant to the research question.

### 2.3 Data Extraction and Analysis

Data extraction was conducted meticulously from various sources within the studies, including text, tables, and figures. Key data points extracted included: authors, year of publication, sample size per group, 16S rRNA primer sequences, DNA extraction kits used, data availability e.g., public repositories, country of origin of the study, inclusion and exclusion criteria, statistical methods employed, mean values of alpha diversity indices, mean values of bacterial abundance, and the choice of Operational Taxonomic Units OTUs versus Amplicon Sequence Variants ASVs for sequence classification.

For studies where data were presented in figures, values were extracted using PlotDigitizer (PlotDigitizer 2024), an image processing software that allows accurate digitization of graphical data. The extracted data were then analyzed using a mean difference test to compare alpha diversity indices and bacterial abundance between diabetic and non-diabetic groups. Subgroup analyses were conducted based on the sequencing method used OTU vs. ASV to explore potential differences in findings related to methodological variations. A random-effects model was employed for meta-analysis, as recommended by Review Manager (Higgins and Green 2011) version 5.4, to account for variability across studies. Statistical significance was determined at a p-value threshold of *<* 0.05. Heterogeneity among studies was assessed using the I^2^ statistic, with the following classifications: low 0%-40%, moderate 30%-60%, substantial 50%-90%, and considerable 75%- 100% (ibid.). All statistical analyses were performed using RStudio (RStudio Team 2024), with R version 4.3.0 (R Core Team 2024) and the ‘meta’ package version 7.0.0 (Schwarzer, Carpenter, and Rücker 2015), ensuring reproducibility and transparency of the analytical process.

## 3 Results

### 3.1 Study Characteristics

The initial search identified 8,410 articles. After applying the inclusion criteria and narrowing down the results, 71 articles were selected for full-text review Figure 1. Following this thorough screening process, thirteen studies met the criteria for inclusion in the final analysis. Of these, nine studies employed Operational Taxonomic Unit OTU sequences, while four utilized Amplicon Sequence Variants ASV for microbiome analysis.

**Figure 1.**
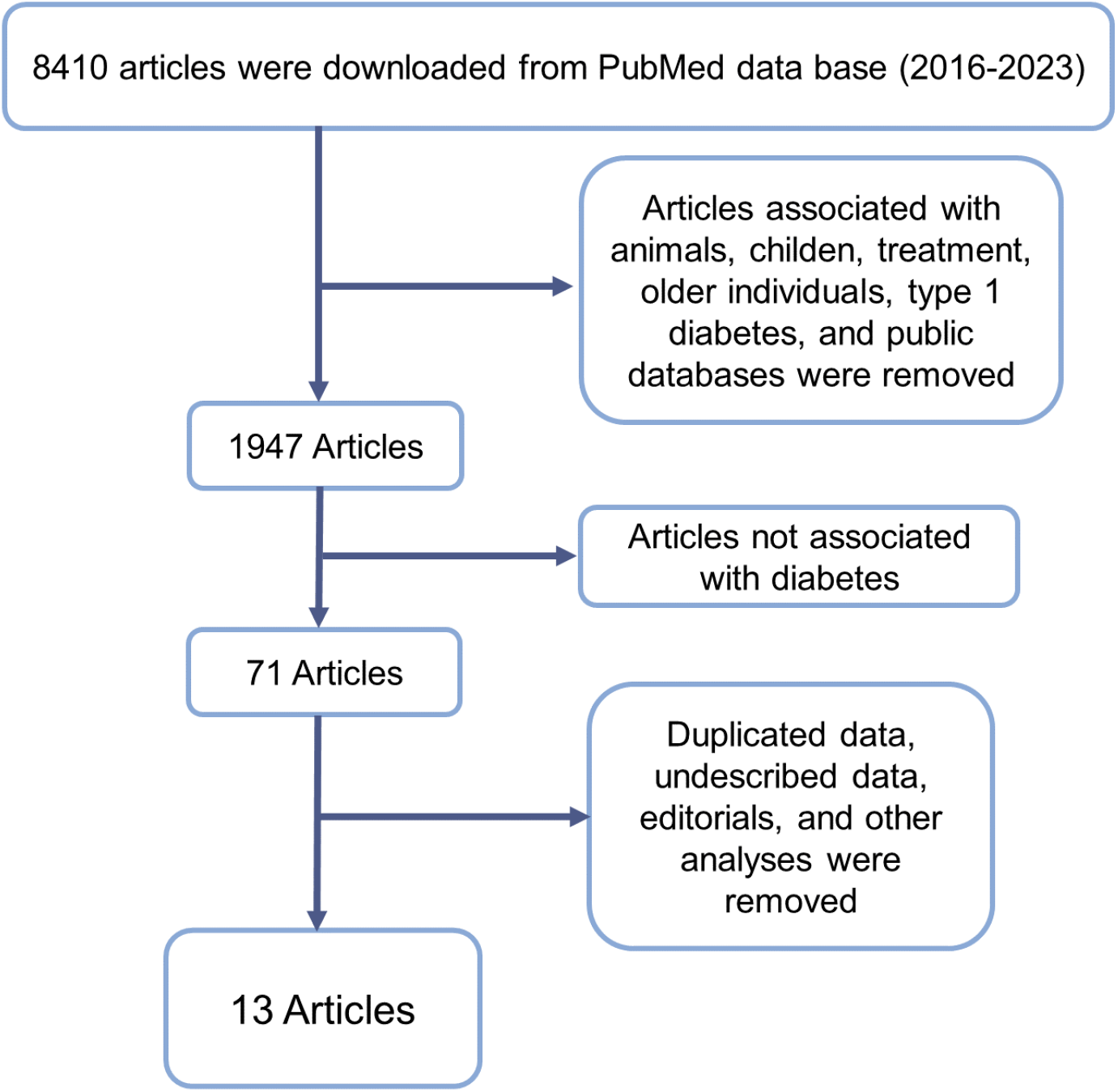
Flowchart representing inclusion and exclusion criteria and resulting article number after exclusion.

Geographically, the majority of the studies were conducted in Asia, with eleven originating from this continent China: 7, Japan: 2, Pakistan: 2. Two studies were conducted in North America USA: 2, and one study was from the Middle East Egypt: 1. Regarding taxonomic classification, the reference databases most frequently used were GreenGenes and SILVA, with each being utilized in four studies. The V3V4 region of the 16S rRNA gene was the most commonly targeted region for primer production, appearing in eight studies Table **??**. Across all included studies, a total of 4,066 sequenced samples were analyzed, providing a robust dataset for the meta-analysis.

**Table I:** Study methodological characteristics.

| Author | Country | Method | 16S Region Reference | Database Reference | N Size |
| --- | --- | --- | --- | --- | --- |
| Ahmad *et al.*, (2019) | Pakistan | OTU | V3V4 | Silva<br>GreenGenes | 60 |
| Ding *et al.*, (2023) | China | ASV | V3V4 | GreenGenes2 | 101 |
| Du *et al.*, (2022) | China | OTU | V3V4 | Not described | 60 |
| Guo *et al.*, (2023) | China | ASV | V3V4 | HOMD | 168 |
| Hashimoto *et al.*, (2020) | Japan | ASV | V3V4 | GreenGenes | 194 |
| Huang *et al.*, (2023) | China | OTU | V4 | GreenGenes | 14 |
| Li *et al.*, (2020) | China | OTU | V4V5 | GreenGenes | 60 |
| Maskarinec *et al.*, (2021) | USA | OTU | V1V3 | Silva | 1702 |
| Salah *et al.*, (2019) | Egypt | OTU | V3V4 | Silva | 60 |
| Saleem *et al.*, (2022) | Pakistan | ASV | V3V4 | Silva | 94 |
| Walker *et al.*, (2021) | USA | OTU | V4 | MetaPhlan2 | 1402 |
| Y. Wang *et al.*, (2017) | China | OTU | V6 | BlastN | 40 |
| J. Wang *et al.*, (2020) | China | OTU | V3V4 | RDB | 171 |

The majority of the studies included in this meta-analysis n = 11 utilized the Shannon index to evaluate alpha diversity between control and diabetic groups. As illustrated in Figure 2, there is substantial heterogeneity among the studies I^2^ *>* 75%, suggesting that multiple factors contribute to the observed variability in alpha diversity results. This high heterogeneity indicates that the results are influenced by differences in study design, population characteristics, sequencing methods, or data analysis techniques.

**Figure 2.**
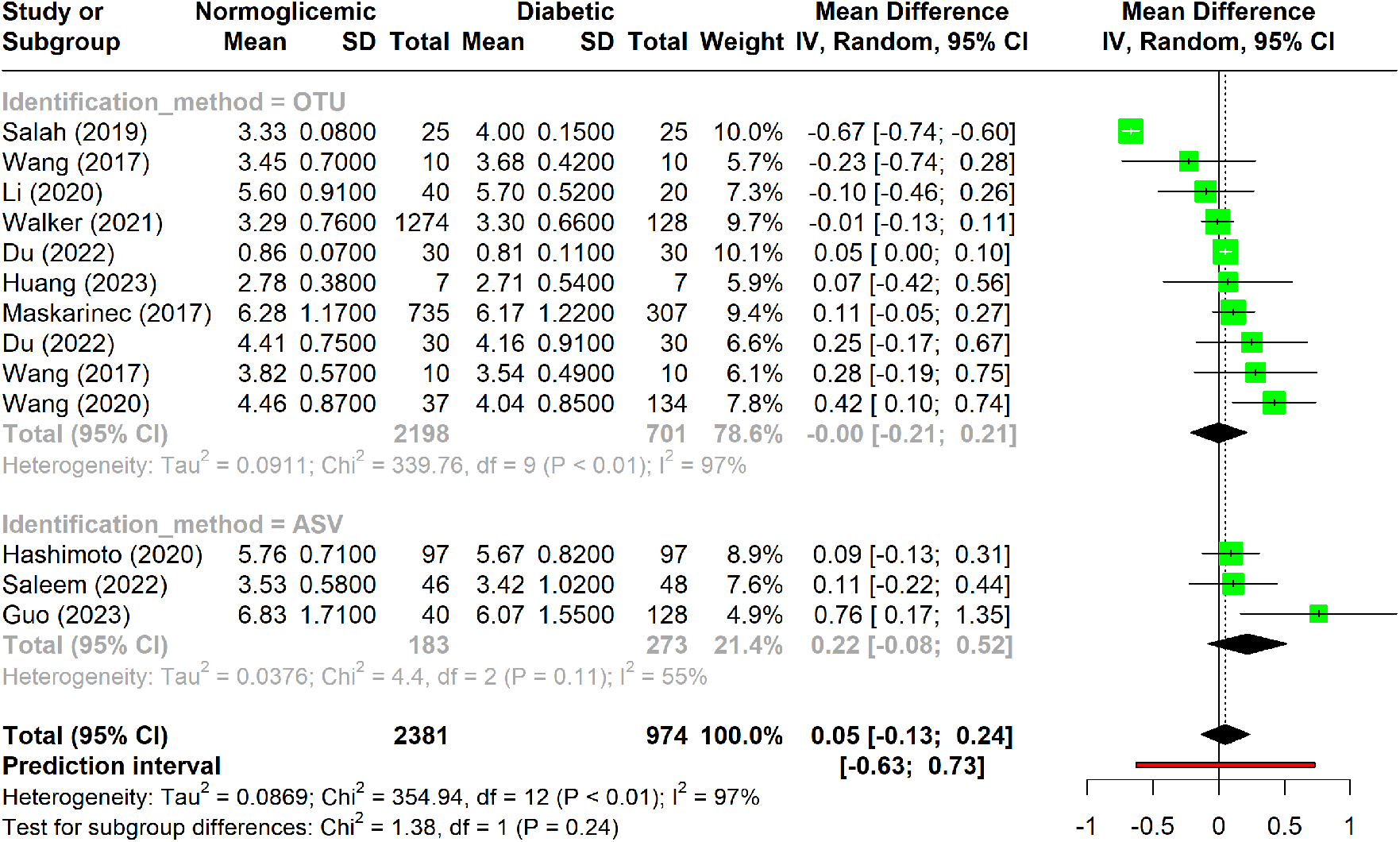
Forest plot of Shannon index in normoglycemic vs. diabetic subjects. Stratified by OTU and ASV identification methods.

Additionally, the results show considerable variation in the Shannon index across studies, as reflected by the wide confidence intervals and the non-significant p-value, which suggests no consistent difference in alpha diversity between diabetic and non-diabetic groups. When the studies were stratified into subgroups based on the identification method, it became evident that studies using the OTU approach exhibited greater heterogeneity compared to those employing the ASV method. The relatively low variability among ASV-based studies could be partly due to the smaller number of studies in this subgroup only three, which may limit the generalizability of these findings.

When the Chao1 index data from all included studies were analyzed using a forest plot, substantial heterogeneity was observed, with I^2^ values falling within the range of 50% to 75% Figure 3. This suggests that while there is notable variability among studies, it is not extreme.

**Figure 3.**
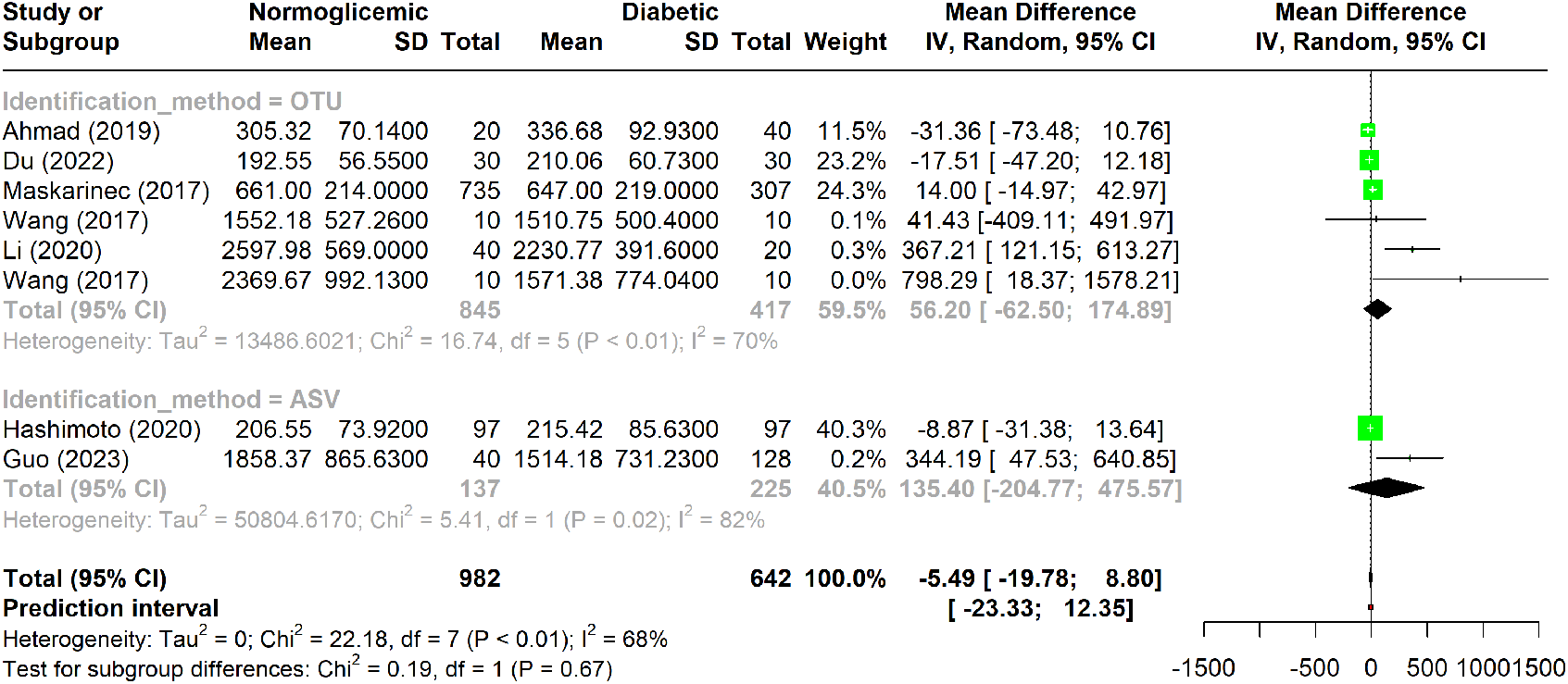
Forest plot of Chao1 index in normoglycemic vs. diabetic subjects. Stratified by OTU and ASV identification methods.

In four studies, higher alpha diversity was reported in diabetic individuals when OTUs were used for analysis. This trend was similarly observed in the ASV data, where two studies indicated an increased Chao1 index in diabetes, although the results were not consistent.

A specific subgroup of three studies that employed the OTU method exhibited substantial heterogeneity I^2^ *<* 75% and showed statistically significant variation p-value *<* 0.01. However, despite this variation, no significant difference was found between the diabetic and non-diabetic groups within this subgroup.

Five studies included in the analysis utilized the phylogenetic whole tree index to evaluate alpha diversity Figure 4. This index showed substantial variation, with heterogeneity ranging from 50% to 90% I^2^, suggesting notable variability across studies. Despite this, the overall p-value was significant p *<* 0.01, indicating that there was no significant difference in phylogenetic distances between diabetic and non-diabetic groups. Among these studies, only one employed the ASV method, while the remaining four used the OTU method. The studies using the OTU method exhibited higher heterogeneity compared to the combined analysis of all five studies. Despite these methodological differences, none of the indices showed a significant difference in alpha diversity between the diabetic and non-diabetic groups.

**Figure 4.**
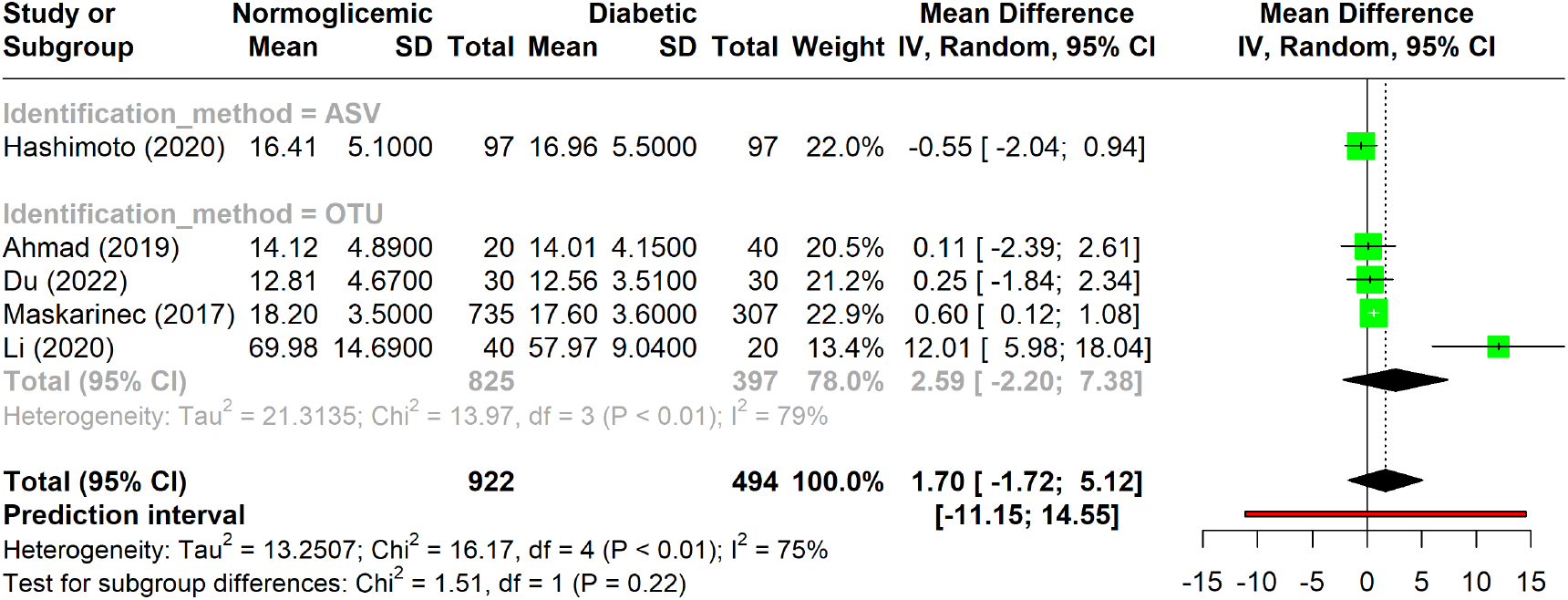
Forest plot of phylogenetic whole tree index in normoglycemic vs diabetic subjects. Stratified by OTU and ASV identification methods.

### 3.2 Taxonomic Composition

The analysis aimed to determine whether diabetic individuals have a distinct abundance of specific phyla compared to non-diabetic individuals. However, no clear trend was observed across the studies. Significant heterogeneity was evident I^2^ *>* 75%, highlighting the diversity in the collected data. This variability suggests that the underlying factors contributing to differences in phylum abundance remain unclear and require further investigation.

Among the 13 studies analyzed, four phyla were frequently associated with diabetes, each showing considerable variation I^2^ *>* 75% and significant p-values p *<* 0.05 Figure 5. This disparity underscores the need for additional research to better understand these associations.

**Figure 5.**
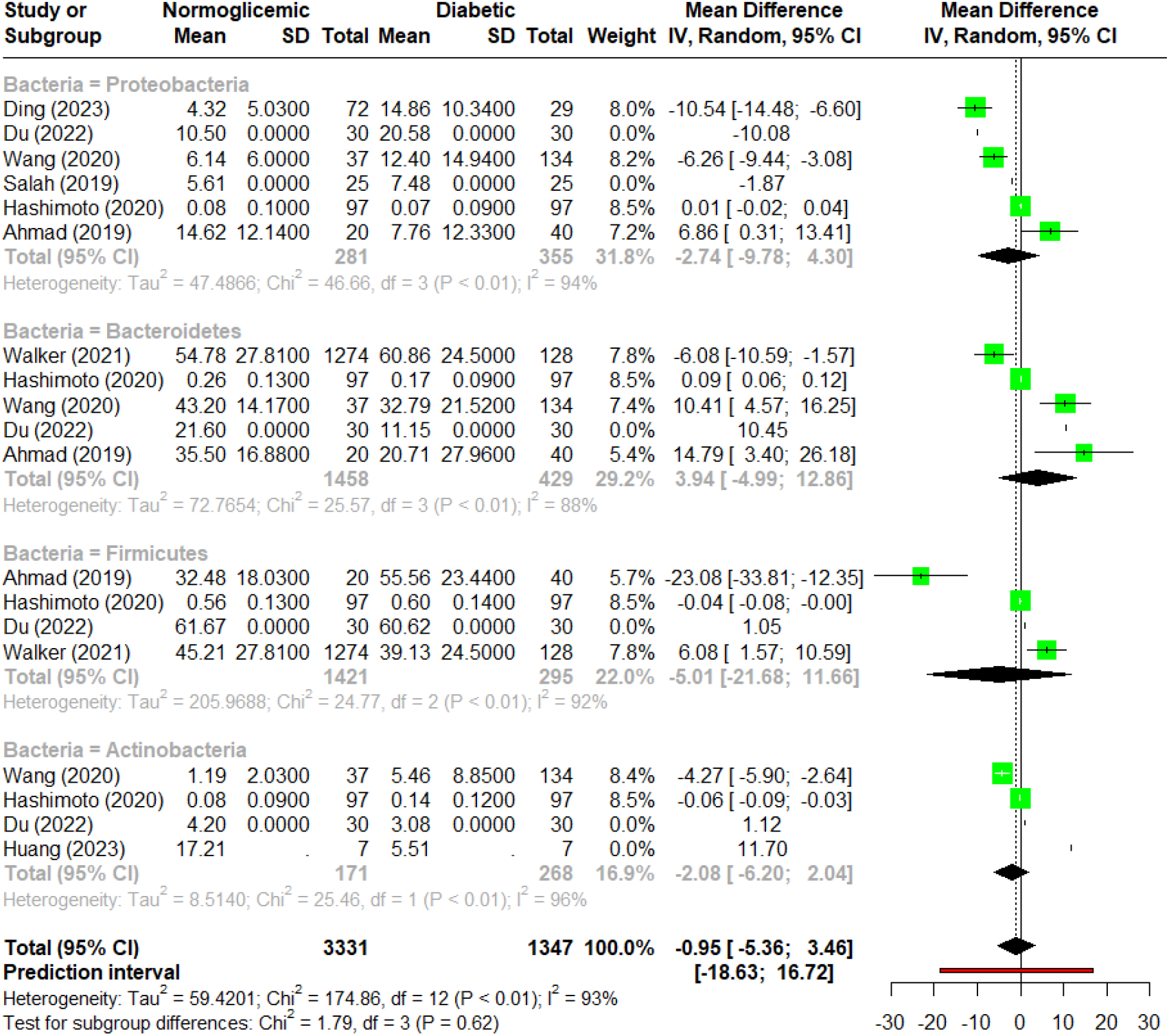
Forest plot of abundance of measurements among associated phyla.

Proteobacteria was the most commonly reported phylum, appearing in six studies. Despite its frequent mention, there was no consensus on its relationship with diabetes. The studies showed high heterogeneity I^2^ = 94%, and no significant differences were found between normoglycemic and diabetic individuals concerning Proteobacteria abundance. Furthermore, factors such as dietary habits and population characteristics, which may influence microbiota composition, have not been thoroughly investigated in this context.

Bacteroidetes was the second most commonly associated phylum, mentioned in five studies. It was the only phylum with heterogeneity below 90%. The findings were mixed: Ahmad *et al*., (2019) and Du *et al*., (2022) reported higher Bacteroidetes abundance in diabetic individuals, while Hashimoto *et al*., (2020) and Walker *et al*., (2021) found lower levels in diabetics. The confidence intervals and p-values suggest that there is no clear association between Bacteroidetes abundance and diabetes.

Firmicutes and Actinobacteria were each associated with diabetes in four studies. Firmicutes exhibited a large confidence interval, mainly due to the findings in Ahmad *et al*., (2019), which indicated a significant difference in abundance between groups. However, the other three studies did not support this result. As for Actinobacteria, although slight variations in means were observed, the p-values and confidence intervals indicate no significant relationship between its abundance and diabetes.

## 4 Discussion

The association between complex traits such as Type 2 Diabetes Mellitus DM2 and gut microbiota has been extensively proposed in the literature (Larsen *et al*., 2010; Baothman *et al*., 2016; Doumatey *et al*., 2020) However, our analysis reveals no significant differences in alpha diversity between normoglycemic and diabetic groups. This outcome is likely influenced by the substantial heterogeneity observed across the studies, suggesting that variations in results may be driven by multiple factors beyond microbial diversity, including methodological differences, personal eating habits, and population characteristics.

A key methodological factor is the choice between OTU and ASV approaches, with most studies favoring OTUs n = 8. The OTU method, while common, is prone to replication issues due to its reliance on clustering algorithms, potentially merging different sequences into the same cluster. On the other hand, the ASV method, particularly when using the DADA2 workflow, offers more precise sequence identification through machine-learning algorithms and stricter merging criteria. Studies have shown that these methodological differences can lead to varying alpha diversity values even when analyzing the same dataset (Joos *et al*., 2020; Chiarello *et al*., 2022). Our results suggest that the lack of significant findings may stem from these methodological disparities, underscoring the need for standardized approaches in microbiome research.

Another critical factor is the sequencing depth, which can significantly impact alpha diversity indices. Indices like Shannon and Simpson’s are relatively robust, but Chao1, which was frequently used in these studies, is more sensitive to sequencing depth variations. This sensitivity might contribute to the observed variability, particularly when comparing OTU and ASV methods (Chiarello *et al*., 2022; Ramakodi 2021). Additionally, the small sample sizes in most studies 10 to 40 individuals per group may not accurately capture the true microbial diversity, introducing another layer of bias.

When evaluating taxonomic composition, our findings indicate no consistent differences in gut microbiota between diabetic and non-diabetic individuals, despite individual studies reporting differential abundances. The choice of reference databases, such as the outdated Greengenes (DeSantis *et al*., 2006; Bolyen *et al*., 2019) or the more recent Silva (Quast *et al*., 2012) can introduce significant variation in taxonomic identification, leading to inconsistent results. This lack of standardization highlights a major challenge in microbiome research, where the diversity of reference databases and methodological approaches creates noise and complicates the interpretation of findings.

The reported alterations in specific phyla, such as Proteobacteria and Bacteroidetes, also exhibit significant heterogeneity I^2^ *>* 75%, suggesting that these findings are not reproducible across studies. For example, Proteobacteria, although frequently associated with diabetes, showed no consistent pattern of alteration, likely due to methodological differences and unconsidered confounding factors such as diet and population-specific characteristics. Similarly, Bacteroidetes, despite being the second most commonly reported phylum, showed conflicting results across studies, further emphasizing the need for standardized methodologies.

The limited statistical power of alpha diversity indices in characterizing gut microbiota is another important consideration. The inherent inter-individual variability in gut microbiome studies necessitates larger sample sizes to achieve reliable assessments (He *et al*., 2018; Kers and Saccenti 2022; Rothschild *et al*., 2018). Most studies analyzed here did not account for this variability adequately, leading to potential biases. The lack of consistent exclusion criteria, such as accounting for recent diarrhea or constipation, can further exacerbate the heterogeneity observed in microbial diversity and abundance (Vandeputte *et al*., 2016; Park *et al*., 2024).

Lifestyle factors, often overlooked in these studies, play a crucial role in shaping the gut microbiome. Recent evidence suggests that microbiota variations are more strongly associated with diet and environmental factors than with disease status alone (He *et al*., 2018; Trefflich *et al*., 2020; Gihawi *et al*., 2023). This perspective aligns with our findings, which indicate that diabetes alone is insufficient to explain the observed microbiota variation. Comprehensive analyses that consider multiple variables are essential for a more accurate understanding of microbiome dynamics.

Finally, the application of 16S rRNA sequencing to human samples presents unique challenges, as even minor environmental differences can lead to significant microbiome variations (Zuniga-Chaves *et al*., 2023). Detailed patient metadata, including dietary habits, stool consistency, and other health conditions, should be a standard inclusion in microbiome studies to improve the reproducibility and interpretability of results. Moreover, integrating metabolic biomarkers with microbiota data may offer more insights into diabetes-related variations than microbiota analysis alone (Yan *et al*., 2023; Gihawi *et al*., 2023).

In conclusion, the reproducibility issues observed in gut microbiota research related to Type 2 diabetes highlight the need for standardized methodologies, comprehensive biological data, and careful consideration of confounding factors. Addressing these challenges is crucial for advancing our understanding of the complex interplay between gut microbiota and metabolic diseases.

In summary, our analysis reveals that the observed inconsistencies across studies on gut microbiota and type 2 diabetes DM2 are likely influenced by methodological differences, particularly in taxonomic identification and reference database selection. To enhance reproducibility in future research, it’s crucial to standardize methodologies and incorporate comprehensive patient metadata, including dietary habits and stool consistency. Additionally, applying advanced statistical techniques, such as bootstrapping, can simulate subpopulations and assess the consistency of findings across these subgroups, offering a more robust understanding of the microbiome’s role in DM2. By addressing these variables and adopting more rigorous statistical approaches, the field can move toward more reliable and reproducible results in microbiome research.

## Acknowledgments

JLPM and GRF concept the idea. JLPM contributed to data acquisition. JLPM and APS wrote the manuscript. JLPM, APS, IM and GRF edited, and draft approved the final version.

## References

Ahmad, Aftab et al., (2019). “Analysis of gut microbiota of obese individuals with type 2 diabetes and healthy individuals”, en. PLOS ONE, Vol. 14 No. 12. Ed. by Jacobs, Jonathan, e0226372. ISSN: 1932-6203. doi: 10.1371/journal.pone.0226372. available at: https://dx.plos.org/10.1371/journal.pone.0226372 (accessed 22 Mar. 2024).

Baothman, Othman A. et al., (2016). “The role of Gut Microbiota in the development of obesity and Diabetes”, en. Lipids in Health and Disease, Vol. 15 No. 1, p. 108. ISSN: 1476-511X. doi: 10.1186/s12944-016-0278-4. available at: http://lipidworld.biomedcentral.com/articles/10.1186/s12944-016-0278-4 (accessed 3 Mar. 2021).

Bolyen, Evan et al., (2019). “Reproducible, interactive, scalable and extensible microbiome data science using QIIME 2”, en. Nature Biotechnology, Vol. 37 No. 8, pp. 852–857. ISSN: 1087-0156, 1546-1696. doi: 10.1038/s41587-019-0209-9. available at: https://www.nature.com/articles/s41587-019-0209-9 (accessed 22 May 2024).

Chiarello, Marlène et al., (2022). “Ranking the biases: The choice of OTUs vs. ASVs in 16S rRNA amplicon data analysis has stronger effects on diversity measures than rarefaction and OTU identity threshold”, en. PLOS ONE, Vol. 17 No. 2. Ed. by Moreno-Hagelsieb, Gabriel, e0264443. ISSN: 1932-6203. doi: 10.1371/journal.pone.0264443. available at: https://dx.plos.org/10.1371/journal.pone.0264443 (accessed 5 Apr. 2024).

DeSantis, T. Z. et al., (2006). “Greengenes, a Chimera-Checked 16S rRNA Gene Database and Workbench Compatible with ARB”, en. Applied and Environmental Microbiology, Vol. 72 No. 7, pp. 5069–5072. ISSN: 0099-2240, 1098-5336. doi: 10.1128/AEM.03006-05.

Ding, Hongying et al., (2023). “Gut microbiome profile of Chinese hypertension patients with and without type 2 diabetes mellitus”, en. BMC Microbiology, Vol. 23 No. 1, p. 254. ISSN: 1471-2180. doi: 10.1186/s12866-023-02967-x. available at: https://bmcmicrobiol.biomedcentral.com/articles/10.1186/s12866-023-02967-x (accessed 22 Mar. 2024).

Doumatey, Ayo P. et al., (2020). “Gut Microbiome Profiles Are Associated With Type 2 Diabetes in Urban Africans”, en. Frontiers in Cellular and Infection Microbiology, Vol. 10, p. 63. ISSN: 2235-2988. doi: 10.3389/fcimb.2020.00063. available at: https://www.frontiersin.org/article/10.3389/fcimb.2020.00063/full (accessed 23 Aug. 2023).

Du, Yage et al., (2022). “Association of gut microbiota with sort-chain fatty acids and inflammatory cytokines in diabetic patients with cognitive impairment: A cross-sectional, non-controlled study”, Frontiers in Nutrition, Vol. 9, p. 930626. ISSN: 2296-861X. doi: 10.3389/fnut.2022.930626. available at: https://www.frontiersin.org/articles/10.3389/fnut.2022.930626/full (accessed 22 Mar. 2024).

Gihawi, Abraham et al., (2023). Major data analysis errors invalidate cancer microbiome findings, en. preprint. Cancer Biology. doi: 10.1101/2023.07.28.550993. available at: http://biorxiv.org/lookup/doi/10.1101/2023.07.28.550993 (accessed 11 Aug. 2023).

Gilbert, Jack A et al., (2018). “Current understanding of the human microbiome”, en. Nature Medicine, Vol. 24 No. 4, pp. 392–400. ISSN: 1078-8956, 1546-170X. doi: 10.1038/nm.4517.

Guo, Xiao-jing et al., (2023). “Distribution characteristics of oral microbiota and its relationship with intestinal microbiota in patients with type 2 diabetes mellitus”, Frontiers in Endocrinology, Vol. 14, p. 1119201. ISSN: 1664-2392. doi: 10.3389/fendo.2023.1119201. available at: https://www.frontiersin.org/articles/10.3389/fendo.2023.1119201/full (accessed 22 Mar. 2024).

Gurung, Manoj et al., (2020). “Role of gut microbiota in type 2 diabetes pathophysiology”, en. EBioMedicine, Vol. 51, p. 102590. ISSN: 23523964. doi: 10.1016/j.ebiom.2019.11.051. available at: https://linkinghub.elsevier.com/retrieve/pii/S235239641930800X (accessed 15 Aug. 2023).

Hashimoto, Yoshitaka et al., (2020). “Intake of sucrose affects gut dysbiosis in patients with type 2 diabetes”, en. Journal of Diabetes Investigation, Vol. 11 No. 6, pp. 1623–1634. ISSN: 2040-1116, 2040-1124. doi: 10.1111/jdi.13293. available at: https://onlinelibrary.wiley.com/doi/10.1111/jdi.13293 (accessed 22 Mar. 2024).

He, Yan et al., (2018). “Regional variation limits applications of healthy gut microbiome reference ranges and disease models”, en. Nature Medicine, Vol. 24 No. 10, pp. 1532–1535. ISSN: 1078-8956, 1546-170X. doi: 10.1038/s41591-018-0164-x. available at: https://www.nature.com/articles/s41591-018-0164-x (accessed 28 July 2023).

Higgins, JPT and Green, S (2011). Cochrane Handbook for Systematic Reviews of Interventions Version 5.1.0 [updated 2022], Accessed: 2024-09-15. The Cochrane Collaboration. available at: https://training.cochrane.org/handbook.

Huang, Sutianzi et al., (2023). “Biotransformation differences of ginsenoside compound K mediated by the gut microbiota from diabetic patients and healthy subjects”, en. Chinese Journal of Natural Medicines, Vol. 21 No. 10, pp. 723–729. ISSN: 18755364. doi: 10.1016/S1875-5364(23)60402-9. available at: https://linkinghub.elsevier.com/retrieve/pii/S1875536423604029 (accessed 22 Mar. 2024).

International Diabetes Federation (2021). IDF Diabetes Atlas, 10th edn, International Diabetes Federation. available at: https://www.diabetesatlas.org.

Joos, Lisa et al., (2020). “Daring to be differential: metabarcoding analysis of soil and plant-related microbial communities using amplicon sequence variants and operational taxonomical units”, en. BMC Genomics, Vol. 21 No. 1, p. 733. ISSN: 1471-2164. doi: 10.1186/s12864-020-07126-4. available at: https://bmcgenomics.biomedcentral.com/articles/10.1186/s12864-020-07126-4 (accessed 6 May 2024).

Kers, Jannigje Gerdien and Saccenti, Edoardo (2022). “The Power of Microbiome Studies: Some Considerations on Which Alpha and Beta Metrics to Use and How to Report Results”, en. Frontiers in Microbiology, Vol. 12, p. 796025. ISSN: 1664-302X. doi: 10.3389/fmicb.2021.796025. available at: https://www.frontiersin.org/articles/10.3389/fmicb.2021.796025/full (accessed 29 Mar. 2024).

Larsen, Nadja et al., (2010). “Gut Microbiota in Human Adults with Type 2 Diabetes Differs from Non-Diabetic Adults”, en. PLoS ONE, Vol. 5 No. 2. Ed. by Bereswill, Stefan, e9085. ISSN: 1932-6203. doi: 10.1371/journal.pone.0009085. available at: https://dx.plos.org/10.1371/journal.pone.0009085 (accessed 19 Aug. 2023).

Li, Qian et al., (2020). “Implication of the gut microbiome composition of type 2 diabetic patients from northern China”, en. Scientific Reports, Vol. 10 No. 1, p. 5450. ISSN: 2045-2322. doi: 10.1038/s41598-020-62224-3. available at: https://www.nature.com/articles/s41598-020-62224-3 (accessed 22 Mar. 2024).

Maskarinec, Gertraud et al., (2021). “The gut microbiome and type 2 diabetes status in the Multiethnic Cohort”, en. PLOS ONE, Vol. 16 No. 6. Ed. by Jacobs, Jonathan, e0250855. ISSN: 1932-6203. doi: 10.1371/journal.pone.0250855. available at: https://dx.plos.org/10.1371/journal.pone.0250855 (accessed 4 Jan. 2024).

MetaHIT consortium et al., (2015). “Disentangling type 2 diabetes and metformin treatment signatures in the human gut microbiota”, en. Nature, Vol. 528 No. 7581, pp. 262–266. ISSN: 0028-0836, 1476-4687. doi: 10.1038/nature15766. available at: https://www.nature.com/articles/nature15766 (accessed 15 Aug. 2023).

Park, Gwoncheol et al., (2024). “Deciphering the Impact of Defecation Frequency on Gut Microbiome Composition and Diversity”, en. International Journal of Molecular Sciences, Vol. 25 No. 9, p. 4657. ISSN: 1422-0067. doi: 10.3390/ijms25094657. available at: https://www.mdpi.com/1422-0067/25/9/4657 (accessed 22 May 2024). PlotDigitizer (2024). PlotDigitizer, https://plotdigitizer.com/app. Accessed: 2024-09-15.

Qin, Junjie et al., (2012). “A metagenome-wide association study of gut microbiota in type 2 diabetes”, en. Nature, Vol. 490 No. 7418, pp. 55–60. ISSN: 0028-0836, 1476-4687. doi: 10.1038/nature11450.

Quast, Christian et al., (2012). “The SILVA ribosomal RNA gene database project: improved data processing and web-based tools”, en. Nucleic Acids Research, Vol. 41 No. D1, pp. D590–D596. ISSN: 0305-1048, 1362-4962. doi: 10.1093/nar/gks1219. available at: http://academic.oup.com/nar/article/41/D1/D590/1069277/The-SILVA-ribosomal-RNA-gene-database-project (accessed 9 July 2024).

R Core Team (2024). R: A Language and Environment for Statistical Computing, Accessed: 2024-09-15. R Foundation for Statistical Computing. Vienna, Austria. available at: https://www.R-project.org.

Ramakodi, Meganathan P. (2021). “Effect of Amplicon Sequencing Depth in Environmental Microbiome Research”, en. Current Microbiology, Vol. 78 No. 3, pp. 1026–1033. ISSN: 0343-8651, 1432-0991. doi: 10.1007/s00284-021-02345-8. available at: http://link.springer.com/10.1007/s00284-021-02345-8 (accessed 7 Apr. 2024).

Rogers, Geraint B and Wesselingh, Steve (2016). “Precision respiratory medicine and the microbiome”, en. The Lancet Respiratory Medicine, Vol. 4 No. 1, pp. 73–82. ISSN: 22132600. doi: 10.1016/S2213-2600(15)00476-2. available at: https://linkinghub.elsevier.com/retrieve/pii/S2213260015004762 (accessed 23 Mar. 2024).

Rothschild, Daphna et al., (2018). “Environment dominates over host genetics in shaping human gut microbiota”, en. Nature, Vol. 555 No. 7695, pp. 210–215. ISSN: 0028-0836, 1476-4687. doi: 10.1038/nature25973. available at: https://www.nature.com/articles/nature25973 (accessed 7 Aug. 2023).

RStudio Team (2024). RStudio: Integrated Development for R, Accessed: 2024-09-15. RStudio, PBC. Boston, MA. available at: http://www.rstudio.com.

Salah, Mohammed et al., (2019). “New Insights on Obesity and Diabetes from Gut Microbiome Alterations in Egyptian Adults”, en. OMICS: A Journal of Integrative Biology, Vol. 23 No. 10, pp. 477–485. ISSN: 1557-8100. doi: 10.1089/omi.2019.0063. available at: https://www.liebertpub.com/doi/10.1089/omi.2019.0063 (accessed 22 Mar. 2024).

Saleem, Afshan et al., (2022). “Unique Pakistani gut microbiota highlights population-specific microbiota signatures of type 2 diabetes mellitus”, en. Gut Microbes, Vol. 14 No. 1, p. 2142009. ISSN: 1949-0976, 1949-0984. doi: 10.1080/19490976.2022.2142009. available at: https://www.tandfonline.com/doi/full/10.1080/19490976.2022.2142009 (accessed 22 Mar. 2024).

Schwarzer, Guido, Carpenter James R., and Rücker, Gerta (2015). Meta-Analysis with R, en. Use R! Springer International Publishing. ISBN: 978-3-319-21415-3 978-3-319-21416-0. doi: 10.1007/978-3-319-21416-0. available at: https://link.springer.com/10.1007/978-3-319-21416-0 (accessed 5 July 2024).

Trefflich, Iris et al., (2020). “Is a vegan or a vegetarian diet associated with the microbiota composition in the gut? Results of a new cross-sectional study and systematic review”, en. Critical Reviews in Food Science and Nutrition, Vol. 60 No. 17, pp. 2990–3004. ISSN: 1040-8398, 1549-7852. doi: 10.1080/10408398.2019.1676697. available at: https://www.tandfonline.com/doi/full/10.1080/10408398.2019.1676697 (accessed 18 Aug. 2023).

Vandeputte, Doris et al., (2016). “Stool consistency is strongly associated with gut microbiota richness and composition, enterotypes and bacterial growth rates”, en. Gut, Vol. 65 No. 1, pp. 57–62. ISSN: 0017-5749, 1468-3288. doi: 10.1136/gutjnl-2015-309618. available at: https://gut.bmj.com/lookup/doi/10.1136/gutjnl-2015-309618 (accessed 15 Mar. 2024).

Vatanen, Tommi et al., (2018). “The human gut microbiome in early-onset type 1 diabetes from the TEDDY study”, en. Nature, Vol. 562 No. 7728, pp. 589–594. ISSN: 0028-0836, 1476-4687. doi: 10.1038/s41586-018-0620-2.

Vrieze, Anne et al., (2012). “Transfer of Intestinal Microbiota From Lean Donors Increases Insulin Sensitivity in Individuals With Metabolic Syndrome”, en. Gastroenterology, Vol. 143 No. 4, 913–916.e7. ISSN: 00165085. doi: 10.1053/j.gastro.2012.06.031. available at: https://linkinghub.elsevier.com/retrieve/pii/S001650851200892X (accessed 9 Aug. 2023).

Walker, Rebecca L. et al., (2021). “Population study of the gut microbiome: associations with diet, lifestyle, and cardiometabolic disease”, en. Genome Medicine, Vol. 13 No. 1, p. 188. ISSN: 1756-994X. doi: 10.1186/s13073-021-01007-5. available at: https://genomemedicine.biomedcentral.com/articles/10.1186/s13073-021-01007-5 (accessed 22 Mar. 2024).

Wang, Jiajia et al., (2020). “Enterotype Bacteroides Is Associated with a High Risk in Patients with Diabetes: A Pilot Study”, en. Journal of Diabetes Research, Vol. 2020, pp. 1–11. ISSN: 2314-6745, 2314-6753. doi: 10.1155/2020/6047145. available at: https://www.hindawi.com/journals/jdr/2020/6047145/ (accessed 22 Mar. 2024).

Wang, Ye et al., (2017). “Gut microbiome analysis of type 2 diabetic patients from the Chinese minority ethnic groups the Uygurs and Kazaks”, en. PLOS ONE, Vol. 12 No. 3. Ed. by Nerurkar Pratibha V., e0172774. ISSN: 1932-6203. doi: 10.1371/journal.pone.0172774. available at: https://dx.plos.org/10.1371/journal.pone.0172774 (accessed 22 Mar. 2024).

WHO (2023). Diabetes-World Health Organization (WHO), Accessed: 2023-04-05. available at: https://www.who.int/news-room/fact-sheets/detail/diabetes.

WHO (2024). Diabetes-World Health Organization (WHO), Accessed: 2024-06-21. available at: https://www.who.int/news-room/fact-sheets/detail/cardiovascular-diseases-(cvds).

Yamaguchi, Yoshiharu et al., (2016). “Association of Intestinal Microbiota with Metabolic Markers and Dietary Habits in Patients with Type 2 Diabetes”, en. Digestion, Vol. 94 No. 2, pp. 66–72. ISSN: 0012-2823, 1421-9867. doi: 10.1159/000447690.

Yan, Min et al., (2023). “Mechanismbased role of the intestinal microbiota in gestational diabetes mellitus: A systematic review and meta-analysis”, Frontiers in Immunology, Vol. 13, p. 1097853. ISSN: 1664-3224. doi: 10.3389/fimmu.2022.1097853. available at: https://www.frontiersin.org/articles/10.3389/fimmu.2022.1097853/full (accessed 22 May 2024).

Zhou, Wenyu et al., (2019). “Longitudinal multi-omics of host–microbe dynamics in prediabetes”, en. Nature, Vol. 569 No. 7758, pp. 663–671. ISSN: 0028-0836, 1476-4687. doi: 10.1038/s41586-019-1236-x.

Zuniga-Chaves, Ibrahim et al., (2023). “Neighborhood socioeconomic status is associated with low diversity gut microbiomes and multi-drug resistant microorganism colonization”, en. npj Biofilms and Microbiomes, Vol. 9 No. 1, pp. 1–9. ISSN: 2055-5008. doi: 10.1038/s41522-023-00430-3. available at: https://www.nature.com/articles/s41522-023-00430-3 (accessed 29 May 2024).

